# Mannose-binding lectin gene polymorphisms in the East Siberia and Russian Arctic populations

**DOI:** 10.1101/2020.05.31.126672

**Authors:** Sergey Yu Tereshchenko, Marina V Smolnikova, Maxim B Freidin

**Author notes:** Corresponding author: Sergey Tereshchenko, Scientific Research Institute of Medical Problems of the North, Partizana Geleznyaka, 3 G, Krasnoyarsk, 660022, Russia. All authors have contributed equally.

## Abstract

**Background:** Mannose-binding lectin (MBL) encoded by *MBL2* gene is a protein with the ability to form carbohydrate complexes with microbial wall promoting their subsequent elimination. Genetically determined levels of MBL can modify the risk and clinical characteristics of many infectious diseases. The frequency of *MBL2* genotypes exhibits significant population differences. The data on the distribution of *MBL2* genotypes among the aborigines of the Russian Arctic territories have not yet been published.

**Methods:** A total of 880 specimens of dried blood spots of the newborns were genotyped. The newborns represented four populations: Nenets, Dolgan-Nganasans, Mixed aboriginal population, and Russians (Caucasians, Krasnoyarsk). Six polymorphisms of the *MBL2* gene were studied: rs11003125, rs7096206, rs7095891, rs5030737, rs1800450, and rs1800451.

**Results:** The frequency of the combined rare O allele (composed of the coding region variants rs5030737, rs1800450, and rs1800451) in the homozygous state was significantly higher in Russians: 10% vs 2% in Nenets and 1% in Dolgan-Nganosans (p<0.001 for Russians vs other populations). The frequency of the high-producing haplotype (HYPA) was 35.4% in the Russian newborns, in keeping with European populations (27-33%); 64% for Nenets and 56% for Dolgan-Nganasans, similar to the estimates obtained for Eskimos and North Amerinds (64-81%).

**Conclusion:** Our study results are in line with the hypothesis that human evolution has been moving in the direction of accumulation of the genotypes associated with low activity of the lectin complement activation pathway because of the prevalence of some intracellular infections such as tuberculosis, whereby low MBL activity may have a protective effect.

## INTRODUCTION

The complement system is a key component of innate immunity, with the main function being predominantly intravascular elimination of bacterial agents. In addition, complement proteins act as a bridge between the innate and adaptive immunity systems, providing adequate conditions for B- and T-lymphocyte maturation and differentiation. The complement system includes both plasma proteins and membrane receptors. The former interacts with each other in three well-known «cascading pathways», i.e. lectin (the most phylogenetically ancient), alternative and classical ones.

Lectins are the general term for proteins forming a separate superfamily for pattern-recognizing receptors with the ability to recognize and aggregate oligo- and polysaccharides. Among all lectins, ficolins (common domain is fibrinogen) and collectins (common domain is collagen) have the unique function to form carbohydrate complexes with microbial wall. These include mannose-binding lectin (MBL), hepatic and renal collectins,[1-4]. The polysaccharide complex of microbial wall with collectin / ficolin and specific proteases leads to the activation of the complement system, the inflammatory reaction and the elimination of bacteria. This activation pathway is called lectin pathway, as opposed to other classical and alternative pathways.

MBL is a conventional C-type lectin comprising several subunits and prone to oligomerization to dimers, trimers and tetramers. The oligomerization properties are genetically determined and critically increase the activity of MBL in terms of the binding of bacterial polysaccharides and complement activation,[1]. Dominant mutations in exon 1 of the *MBL2* gene, located on chromosome 10 (10q21.1), have been known to result in an impaired ability of MBL to oligomerize and, accordingly, to the reduced plasma concentration and functional activity. Mutations in codons 52 (rs5030737; A / D), 54 (rs1800450; A / B) and 57 (rs1800451; A / C) lead to the similar consequences. Alleles containing mutations in codons 52, 54 and 57 are designated as D, B and C, respectively, as contrasted with the wild-type allele (A). Mutations D, B and C are commonly combined as the collective designation “O” due to the physiological consequences of the same type.

In addition to the coding mutations in exon 1, the immunological function of MBL is also influenced by the gene promoter mutations: dimorphisms at the rs11003125 (H / L) and rs7096206 (Y / X) loci modulate transcriptional activity, thereby significantly affecting the concentration of MBL in blood plasma (H> L and Y> X),[1]. The HY haplotype has been found to be associated with the highest plasma concentration of MBL, the LY haplotype is associated with an average concentration, and the LX is associated with the low concentration,[5]. Besides, dimorphism in the non-coding region of exon 1 (rs7095891; P / Q) has been found.

Due to the pronounced linkage disequilibrium, all the reported mutations can be combined into a limited number of haplotypes (HYPA, LXPA, LYQA, LYPA, HYPD, LYPB, LYPD and LYQC) out of 64 possible,[5, 6]. The frequency of *MBL2* haplotypes has significant population differences,[5, 7]. Thus, the HYPA haplotype frequency associated with the high MBL concentration varies from 6-8% in African populations (Mozambique, Kenya,[5, 8]) to 64-81% in northern indigenous populations (Native North-Americans, Inuits,[9-11]). In this gradation, Caucasians, with 27-30% frequency of the HYPA haplotype, occupy an intermediate position,[12-14].

Additionally, to assess the clinical consequences of genetically determined differences in MBL expression, MBL-deficient (YO / YO or XA / YO), MBL-intermediate (YA / YO or XA / XA) and MBL-highly expressing (YA / YA or XA / YA) diplotypes were proposed to be distinguished,[10, 15, 16]. It is generally believed that 20-25% of the entire human population are MBL-deficient haplotype carriers, and there is no or extremely low MBL concentration in plasma in 8-10%,[5, 17, 18].

The vast majority of MBL-deficient individuals are generally healthy. There are evident clinical consequences for MBL deficiency only in certain clinical situations, i.e. in patients with neutropenia, after organ and tissue transplantation, in newborns, especially in premature infants,[19, 20]. Nevertheless, a large number of studies have shown that genetically determined levels of MBL can modify the risk and clinical characteristics of many infectious diseases. Moreover, this effect seems to be pluripotent in nature. A sufficiently high MBL level is a protective factor against the occurrence and severity of infections caused by encapsulated bacteria (*Streptococcus pneumoniae, Haemophilus influenzae, and Neisseria meningitidis*), primarily in young children,[21, 22]. At the same time, it was hypothesized that normal / high MBL levels may increase the risk of infection and hyperinflammatory response in disorders caused by certain intracellular pathogens such as *Mycobacterium tuberculosis* and *Leishmania*,[18, 23]. Therefore, the carriers of certain MBL-deficient haplotypes may have a specific clinical advantage with the intracellular infections. The recent meta-analyses have shown that the relationship between MBL genotypes and tuberculosis are controversial, i.e. some genetic variations increase the risk of the disease (rs1800450, rs5030737), while some may decrease (rs1800451, rs7095891),[24-26]. The analysis is complicated by large heterogeneity in different clinical forms of tuberculosis in the studies. In addition, risk assessment can largely depend on the ethnicity and age of the populations,[24, 25, 27].

To the best of our knowledge, the data on the prevalence of genotypes and haplotypes of the *MBL2* gene in the Russian population of East Siberia and among the aborigines of the Russian Arctic territories have not yet been published. The current study aims to fill this gap by the analysis of the aboriginal and alien populations of this region: the Nenets, the Dolgans-Nganasans and the Russians.

## METHODS

A total of 880 specimens of dried blood spots for the newborns obtained from the Krasnoyarsk Regional Consulting-Diagnostic Centre for Medical Genetics to study the prevalence of single nucleotide polymorphisms of *MBL2* gene.

The demographic characteristics of the studied newborns according to the region of mother’s settlement was the same as in our previously published study,[28]. The newborns were split into four groups to study ethnic specificity of the *MBL2* polymorphisms: (1) 260 from the Arctic region of mother’s settlement, from villages with predominantly Nenets population (Nenets comprise 85% of the population); (2) 110 from the Arctic region of mother’s settlement, from villages with predominantly Dolgan-Nganasan population (Dolgan-Nganasans comprise 91% of the population); (3) 210 from the Arctic region of mother’s settlement, from villages with mixed populations with various combination of indigenous and alien residents; (4) 300 newborns of European ancestry (Russians by self-reports of their mothers) from the city of Krasnoyarsk.

The study was approved by the Ethical Committee of the Scientific Research Institute of Medical Problems of the North (# 9 of 8.09.2014). Signed informed consent was obtained from parents of all participated children.

### Blood Sample Collection and Genotyping

DNA was extracted using DIAtomTM DNA Prep kits (“Izogen”, Russia). Genotyping was carried out using restriction fragment lengths polymorphism approach (RFLP) and real-time polymerase chain reaction. Six polymorphisms of the *MBL2* gene were studied: rs11003125, rs7096206, rs7095891, rs5030737, rs1800450 and rs1800451.

Genotyping of rs1800450 and rs1800451 polymorphisms was performed by RFLP approach. The relevant genomic fragment of 349 bp was amplified using the pair of oligonucleotide primers: forward 5’-TAGGACAGAGGGCATGCTC-3’ and reverse 5’-CAGGCAGTTTCCTCTGGAAGG-3’ (annealing temperature 60°C). Restriction endonucleases *AccB1 I* (rs1800450) and *Mbo II* (rs1800451) for hydrolysis of the fragment followed by the electrophoresis in agarose gel with ethidium bromide to visualise the results. For rs1800450, *AccB1 I* endonuclease produces a single fragment of 349 bp for the B allele and two fragments of 260 and 89 bp for the A allele. For rs1800451, *Mbo II* endonuclease produces a single fragment of 349 bp for the A allele and two fragments of 270 and 79 bp for the C allele.

Four polymorphisms rs11003125, rs7096206, rs7095891 and rs5030737 were carried out using the real-time polymerase chain reaction approach. The nucleotide sequences of allele-specific probes are presented in Table 1.

**Table 1.**
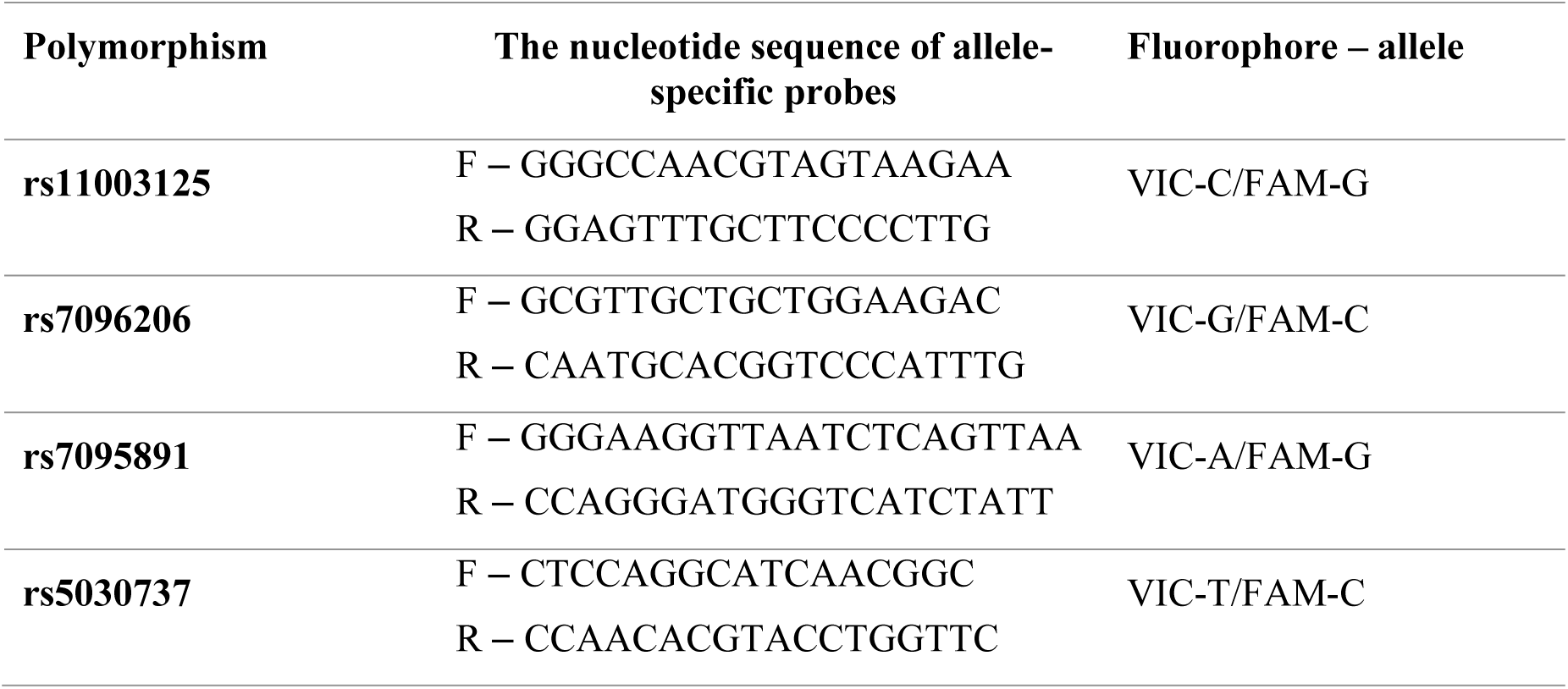
The nucleotide sequence of allele-specific probes used for genotyping

### Statistical Analysis

Differences in genotypic frequencies between the ethnic groups were assessed using the Pearson’s χ2 test. Haplotypes were assessed and compared between the populations using the *haplo*.*stats* package for R. Haplotype score test was applied with 1000 permutations to calculate p-values. Bonferroni correction for multiple testing was duly applied. Statistically significant differences were considered at p<0.05 after correction for multiple testing.

## RESULTS

The genotype frequencies for the polymorphic regions of the MBL2 gene included in the study, except for rs1800451, are presented in Table 2. The variant C allele in rs1800451 was revealed in only one case out of 880 tested newborns, namely in the homozygous state (CC) in a European living in Krasnoyarsk, so this SNP was excluded from subsequent analysis. Among the homozygous variants of the studied *MBL2* gene polymorphisms, the most straightforward population differences were identified for rs11003125 in the promoter region: the frequency of the LL genotype associated with low MBL production in the Russian population was 2-3 times higher than the frequency in the native populations of the Arctic. Namely, in Russians it was 37%, in Nenets it was 10%, in Dolgan-Nganosans it was 15% (p<0.001 for Russians vs other populations). The frequency of the combined rare O allele (composed of the coding region variants rs5030737, rs1800450 and rs1800451) in the homozygous state was also significantly higher in Russians: 10% vs 2% in Nenets and 1% in Dolgan-Nganosans (p<0.001 for Russians vs other populations).

**Table 2.** *MBL2* genotypes frequencies among newborns from different ethnic populations of Taymyr-Dolgan-Nenets region of Krasnoyarskiy Kray and the city of Krasnoyarsk, n (%)

|  |  | Nenets | Dolgans-Nganasans | Mixed Arctic<br>populations | Russians |
| --- | --- | --- | --- | --- | --- |
| MBL2 genotype |  | (n=260) | (n=110) | (n=210) | (n=300) |
| **rs11003125**<br>**promoter** | HH | 114 (0.44) | 32 (0.29) | 71 (0.34) | 54 (0.18) |
|  | HL | 121 (0.47) | 61 (0.55) | 103 (0.49) | 134 (0.45) |
|  | LL | 25 (0.10) | 17 (0.15) | 36 (0.17) | 112 (0.37) |
| **rs7096206**<br>**promoter** | XX | 4 (0.02) | 3 (0.03) | 4 (0.02) | 11 (0.04) |
|  | XY | 60 (0.23) | 33 (0.30) | 47 (0.22) | 115 (0.38) |
|  | YY | 196 (0.75) | 74 (0.67) | 159 (0.76) | 174 (0.58) |
| **rs7095891**<br>**5'UTR** | PP | 4 (0.02) | 3 (0.03) | 4 (0.02) | 11 (0.04) |
|  | PQ | 60 (0.23) | 33 (0.30) | 47 (0.22) | 115 (0.38) |
|  | QQ | 196 (0.75) | 74 (0.67) | 159 (0.76) | 174 (0.58) |
| **rs5030737**<br>**exon 1** | AA | 252 (0.97) | 110 (1.00) | 201 (0.96) | 265 (0.88) |
|  | AD | 8 (0.03) | 0 (0.00) | 9 (0.04) | 32 (0.11) |
|  | DD | 0 (0.00) | 0 (0.00) | 0 (0.00) | 3 (0.01) |
| **rs1800450**<br>**exon 1** | AA | 218 (0.84) | 85 (0.77) | 164 (0.78) | 221 (0.74) |
|  | AB | 37 (0.14) | 24 (0.22) | 35 (0.17) | 54 (0.18) |
|  | BB | 5 (0.02) | 1 (0.01) | 11 (0.05) | 25 (0.08) |
| **Combined**<br>**coding**<br>**genotype**<br>**(rs5030737,**<br>**rs1800450,**<br>**rs1800451)** | AA | 221 (0.81) | 85 (0.77) | 155 (0.74) | 189 (0.63) |
|  | AO | 44 (0.17) | 24 (0.22) | 44 (0.21) | 82 (0.27) |
|  | OO | 5 (0.02) | 1 (0.01) | 11 (0.05) | 29 (0.10) |

The haplotype frequencies for *MBL2* are presented in Table 3. The HYPA haplotype frequency was 35.4% in Russian newborns from East Siberia, similar to the frequencies of European populations (the Netherlands, 27%,[12], Denmark, 30%,[14], Czech Republic, 33%,[13]), as well as Brazilian Caucasians (28-34%,[7, 29]). However, the HYPA haplotype frequency in newborns of the Arctic populations was statistically significantly higher than in the Russians: 64% for the Nenets and 56% for the Dolgan-Nganosans, close to the frequencies identified for the Eskimos (81%,[5, 9]) and the North Amerinds (64%,[11]). At the same time, low frequencies of the MBL-deficient LXPA haplotype were recorded in the Russian newborns (Table 3). The greatest differences in the frequencies of these haplotypes were typical for the Nenets population, with a statistically significant decrease in the LYPA haplotype frequency identified only in this Arctic population.

**Table 3.** *MBL2* haplotypes frequencies among newborns from different ethnic populations of Taymyr-Dolgan-Nenets region of Krasnoyarskiy Kray and the city of Krasnoyarsk

| Population | n | *MBL2* haplotypes |  |  |  |  |  |  |  |
| --- | --- | --- | --- | --- | --- | --- | --- | --- | --- |
|  |  | *HYPA* | *LXPA* | *LYQA* | *LYPA* | *LYPB* | *LYQB* | HYPD | LYPD |
| **Nenets (1)** | 260 | 0.638 | 0.127 | 0.100 | 0.026 | 0.070 | 0.007 | 0.015 | 0 |
| **Dolgans-**<br>**Nganasans (2)** | 110 | 0.556 | 0.154 | 0.118 | 0.033 | 0.116 | 0 | 0 | 0 |
| **Mixed Arctic**<br>**population (3)** | 210 | 0.551 | 0.118 | 0.120 | 0.044 | 0.100 | 0.025 | 0.015 | 0 |
| **Russians (4)** | 300 | 0.354 | 0.221 | 0.133 | 0.048 | 0.145 | 0.025 | 0.045 | 0.018 |
| **p** |  | p1-4<0.001<br>p2-4<0.001<br>p3-4<0.001<br>p1-3=0.036 | p1-4<0.001<br>p3-4<0.001 |  | p1-4<0.001 |  |  | p1-4=0.030<br>p2-4=0.012 |  |

Note: only p-values <0.05 after correction for multiple testing are provided.

Data on the MBL-deficient haplotypes frequencies in the studied populations are summarized in Table 4. MBL-deficient (YO / YO or XA / YO), MBL-intermediate (YA / YO or XA / XA) and MBL-highly expressing (YA / YA or XA / YA) haplotypes were identified. Populations of Nenets and Dolgan-Nganosans showed significantly lower frequencies of MBL-deficient genotypes than Caucasians of East Siberia (3.9%, 6.4% and 21.3%, respectively, p <0.001). The mixed Arctic population has demonstrated an intermediate frequency value of 9.1%.

**Table 4.** The prevalence of the *MBL2* deficient haplotypes among newborns from different ethnic populations of Taymyr-Dolgan-Nenets region of Krasnoyarskiy Kray and the city of Krasnoyarsk, n (%)

| *MBL2* genotypes | Nenets<br>(n=260) | Dolgans-Nganasans<br>(n=110) | Mixed<br>aboriginal<br>population<br>(n=210) | Russians,<br>Krasnoyarsk<br>(n=300) | p |
| --- | --- | --- | --- | --- | --- |
|  | 1 | 2 | 3 | 4 |  |
| ***MBL2* deficient**<br>**haplotype** | 10<br>(3.9%) | 7<br>(6.4%) | 19<br>(9.1%) | 64<br>(21.3%) | 1,2,3-<br>4<0.001<br>1-3=0.02 |
| ***MBL2***<br>**intermediate**<br>**haplotype** | 43<br>(16.5%) | 21<br>(19.1%) | 39<br>(18.6%) | 58<br>(19.3%) |  |
| ***MBL2* sufficient**<br>**haplotype** | 207<br>(79.6%) | 82<br>(74.6%) | 152<br>(72.4%) | 178<br>(59.3%) | 1-4<0.001<br>2-4=0.005<br>3-4=0.002 |

Note: only p-values <0.05 are provided.

## DISCUSSION

It has been suggested that, at the population level, the clinical consequences of an inherently high ability to produce functionally active MBL in the Arctic representatives are low risk of severe bacterial infections at early age and, likely the higher risk of tuberculosis at older age,[18, 26]. Moreover, the low incidence of atherosclerosis and cardio-vascular diseases among the aborigines of the Arctic, along with factors such as high consumption of omega-3 fatty acids and lifestyle features, might be due to the genetic characteristics of the production and activity of MBL. The probability of such a relationship has been described in a number of publications,[9-11, 30].

In the current study, the data on the frequencies of genotypes and haplotypes of the *MBL2* gene among indigenous populations of the Russian Arctic territories have been obtained for the first time. Previously, we showed a high prevalence of the genotypes associated with a high L-ficolin activity in the Arctic populations of Nenets and Dolgan-Nganosans, compared to Caucasians of East Siberia,[28]. In the aboriginal populations of both Nenets and Dolgans-Nganasans, we found the decreased prevalence of the genotype for the rs7851696 polymorphism associated with low L-ficolin carbohydrates binding capacity, as compared to the Russian population. Newborns in mixed arctic populations were characterized by the intermediate prevalence of the rs7851696 rare allele genotype. We concluded that Arctic populations are characterized by a genetic predisposition to the higher level of L-ficolin functional activity, as compared to the Russian population. The Nenets population exhibited several important features as compared with the Dolgans-Nganasans: lower prevalence of the allele T for the rs17549193 polymorphism and higher prevalence of the allele T for the rs7851696 polymorphism. We believe that this genotype is a genetic marker of high functional capacity of L-ficolin in Nenets population. Notably, in the current study Nenets population exhibited slightly higher prevalence MBL sufficient genotypes in comparison to Dolgans-Nganassans (Tables 2, 3).

Thus, the indigenous Arctic populations are genetically characterized by the greater activity of at least two different components of the lectin complement activation pathway, i.e. MBL and L-ficolin. A definite advantage of our approach for estimating population prevalence of MBL- and L-ficolin genotypes is to study the newborns when the probable elimination of unfavorable genetic variations, possible at an older age, has not yet occurred.

Our study results are in line with the hypothesis that human evolution has been moving towards the accumulation of the genotypes associated with low activity of the lectin complement activation pathway because of the prevalence of some intracellular infections such as tuberculosis and leprosy, whereby low MBL and L-ficolin activity may have a protective effect,[7, 18, 23]. Isolated Arctic populations have been suggested to encounter these infections historically later and, therefore, to maintain the high activity of lectin complement activation pathway formed in the early stages of human evolution.

## ACKNOWLEDGEMENTS

We wish to thank “Krasnoyarsk Regional Consulting-Diagnostic Centre for Medical Genetics” staff for helping to collect the samples of dried blood spots.

## CONTRIBUTORS

ST designed and coordinated the study. MS performed the genotyping. ST, MS, MF performed the data analysis and drafted the manuscript. All authors commented on the manuscript and agreed on the final version to be published.

## PATIENT CONSENT

Obtained

## ETHICS APPROVAL

The study was approved by the Ethical Committee of the Scientific Research Institute of Medical Problems of the North (# 9 of 8.09.2014).

